# Rapid and Interpretable Protein Contact Map Prediction Using a Pattern-Matching Strategy

**DOI:** 10.1101/2025.09.08.674800

**Authors:** Aysima Hacisuleyman, Dirk Fasshauer

## Abstract

Protein sequence determines the structure, function, and dynamics of a protein. In recent years, enormous progress has been made in translating sequence information into structural information using machine learning approaches. However, because of the underlying methodology, it is an immense computational challenge to extract this information from the ever-increasing number of sequences. In the present study, we show that it is possible to create two-dimensional contact maps from sequences, for which only a few exemplary structures are available on a laptop without the need for GPUs or high-performance computing clusters. This is achieved by using a pattern matching approach. The resulting contact maps largely reflect the interactions in the three-dimensional structures. The validity of our method was tested on the 25 protein domains, with abundant structural data, achieving correlations of 0.73-0.94 between predicted and experimental contact maps. To demonstrate broader applicability, we further validated our approach on 7,599 poorly annotated sequences using homologous structural templates, achieving a mean F1-score of 0.609 ± 0.095 and mean accuracy of 0.954 ± 0.036 when compared against high-confidence AlphaFold structures. These results demonstrate that our pattern matching approach maintains robust performance even when relying on a small number of structural templates.

## Introduction

Despite the rapid growth of sequenced proteins, experimentally determined structures remain available for only a small fraction, creating a gap between sequence data and structural knowledge. This imbalance limits our ability to infer function, understand molecular mechanisms, and explore protein dynamics at scale. Machine learning-based structure prediction methods, including AlphaFold2^1^, RoseTTAFold^2^ and ESMfold^3^, have transformed structural biology, yet they require substantial computational resources that limit their application in high-throughput analyses.

Contact maps, representing pairwise spatial relationships between amino acid residues, provide a simplified but informative representation of protein structure. These two-dimensional matrices capture essential structural features while reducing computational complexity. Here, we present a pattern-matching approach that leverages existing structural repositories to rapidly generate contact maps for query sequences. Rather than predicting full three-dimensional structures, the method identifies conserved contact patterns from homologous proteins and maps them onto query sequences. By integrating multiple structural templates, including alternative conformations, our approach captures dynamic features of proteins and can identify conserved motifs even when sequence similarity is limited. By analyzing contact patterns across multiple conformational states, one can gain insights into protein dynamics that might otherwise require extensive molecular dynamics simulations.

This strategy offers several advantages: it reduces computational cost, enables high-throughput analyses, and provides biologically meaningful predictions for proteins with limited structural information. To benchmark the method, we analyzed 25 well-characterized protein domains, comparing their predicted contact maps with experimentally determined reference structures. We further evaluated its applicability on poorly annotated sequences lacking comprehensive structural coverage, demonstrating that evolutionarily related homologs can still provide informative patterns for contact prediction.

## Methods

### Sequence and Structure Retrieval

We prepared two complementary datasets for method evaluation. The primary benchmark consisted of 25 well-characterized protein domains from InterPro^4^ (Table 1). We retrieved all available sequences and curated PDB structures for each entry to ensure high-quality annotation and capture conformational diversity. The second dataset comprised of 7,599 poorly annotated proteins from UniProt’s^5^ TrEMBL^6^ collection (Table 2, Table S2). Unreviewed sequences were filtered for low annotation scores(annotation scores of 1), sequence diversity, and high-confidence predicted structures (with average pLDDT ≥ 80). JackHMMER^7^ was used to identify homologous PDB structures, applying additional filters to exclude sequences with excessive (>1,500) or insufficient (<50) structural hits. After filtering, the dataset included proteins with diverse secondary structure compositions representative of the general protein population. Specifics of the selection process are explained and shown in the Zenodo repository.

**Table 1.**
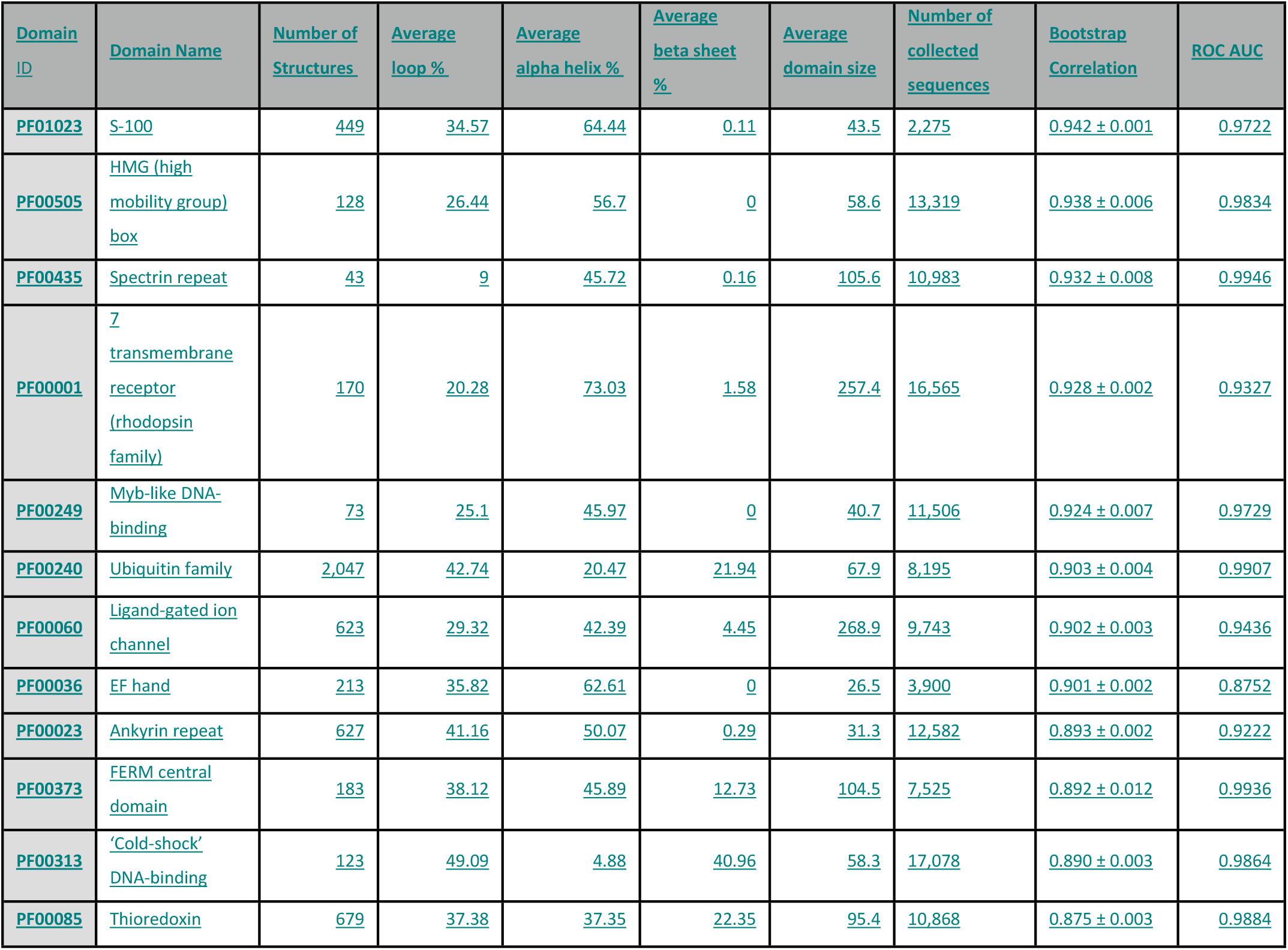
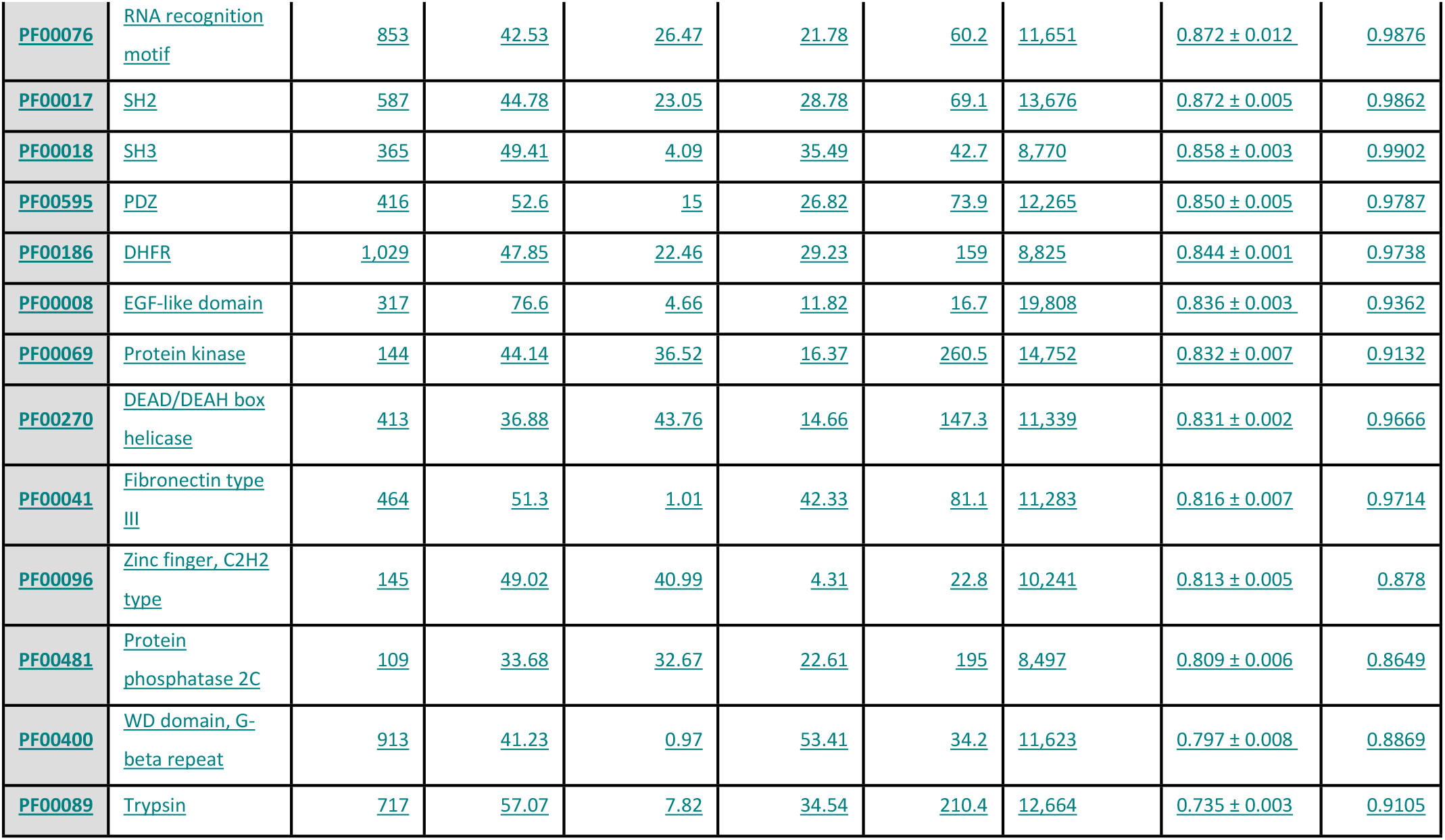
Structural composition and contact prediction performance for 25 well-characterized protein domains

| Domain ID | Domain Name | Number of Structures | Average loop % | Average alpha helix % | Average beta sheet % | Average domain size | Number of collected sequences | Bootstrap Correlation | ROC AUC |
| --- | --- | --- | --- | --- | --- | --- | --- | --- | --- |
| PF01023 | S-100 | 449 | 34.57 | 64.44 | 0.11 | 43.5 | 2,275 | 0.942 ± 0.001 | 0.9722 |
| PF00505 | HMG (high mobility group) box | 128 | 26.44 | 56.7 | 0 | 58.6 | 13,319 | 0.938 ± 0.006 | 0.9834 |
| PF00435 | Spectrin repeat | 43 | 9 | 45.72 | 0.16 | 105.6 | 10,983 | 0.932 ± 0.008 | 0.9946 |
| PF00001 | 7 transmembrane receptor (rhodopsin family) | 170 | 20.28 | 73.03 | 1.58 | 257.4 | 16,565 | 0.928 ± 0.002 | 0.9327 |
| PF00249 | Myb-like DNA-binding | 73 | 25.1 | 45.97 | 0 | 40.7 | 11,506 | 0.924 ± 0.007 | 0.9729 |
| PF00240 | Ubiquitin family | 2,047 | 42.74 | 20.47 | 21.94 | 67.9 | 8,195 | 0.903 ± 0.004 | 0.9907 |
| PF00060 | Ligand-gated ion channel | 623 | 29.32 | 42.39 | 4.45 | 268.9 | 9,743 | 0.902 ± 0.003 | 0.9436 |
| PF00036 | EF hand | 213 | 35.82 | 62.61 | 0 | 26.5 | 3,900 | 0.901 ± 0.002 | 0.8752 |
| PF00023 | Ankyrin repeat | 627 | 41.16 | 50.07 | 0.29 | 31.3 | 12,582 | 0.893 ± 0.002 | 0.9222 |
| PF00373 | FERM central domain | 183 | 38.12 | 45.89 | 12.73 | 104.5 | 7,525 | 0.892 ± 0.012 | 0.9936 |
| PF00313 | ‘Cold-shock’ DNA-binding | 123 | 49.09 | 4.88 | 40.96 | 58.3 | 17,078 | 0.890 ± 0.003 | 0.9864 |
| PF00085 | Thioredoxin | 679 | 37.38 | 37.35 | 22.35 | 95.4 | 10,868 | 0.875 ± 0.003 | 0.9884 |
| PF00076 | RNA recognition motif | 853 | 42.53 | 26.47 | 21.78 | 60.2 | 11,651 | 0.872 ± 0.012 | 0.9876 |
| PF00017 | SH2 | 587 | 44.78 | 23.05 | 28.78 | 69.1 | 13,676 | 0.872 ± 0.005 | 0.9862 |
| PF00018 | SH3 | 365 | 49.41 | 4.09 | 35.49 | 42.7 | 8,770 | 0.858 ± 0.003 | 0.9902 |
| PF00595 | PDZ | 416 | 52.6 | 15 | 26.82 | 73.9 | 12,265 | 0.850 ± 0.005 | 0.9787 |
| PF00186 | DHFR | 1,029 | 47.85 | 22.46 | 29.23 | 159 | 8,825 | 0.844 ± 0.001 | 0.9738 |
| PF00008 | EGF-like domain | 317 | 76.6 | 4.66 | 11.82 | 16.7 | 19,808 | 0.836 ± 0.003 | 0.9362 |
| PF00069 | Protein kinase | 144 | 44.14 | 36.52 | 16.37 | 260.5 | 14,752 | 0.832 ± 0.007 | 0.9132 |
| PF00270 | DEAD/DEAH box helicase | 413 | 36.88 | 43.76 | 14.66 | 147.3 | 11,339 | 0.831 ± 0.002 | 0.9666 |
| PF00041 | Fibronectin type III | 464 | 51.3 | 1.01 | 42.33 | 81.1 | 11,283 | 0.816 ± 0.007 | 0.9714 |
| PF00096 | Zinc finger, C2H2 type | 145 | 49.02 | 40.99 | 4.31 | 22.8 | 10,241 | 0.813 ± 0.005 | 0.878 |
| PF00481 | Protein phosphatase 2C | 109 | 33.68 | 32.67 | 22.61 | 195 | 8,497 | 0.809 ± 0.006 | 0.8649 |
| PF00400 | WD domain, G-beta repeat | 913 | 41.23 | 0.97 | 53.41 | 34.2 | 11,623 | 0.797 ± 0.008 | 0.8869 |
| PF00089 | Trypsin | 717 | 57.07 | 7.82 | 34.54 | 210.4 | 12,664 | 0.735 ± 0.003 | 0.9105 |

**Table 2.** Structural composition and prediction performance for poorly annotated sequences

|  | Value |
| --- | --- |
| Number of sequences analysed | 7,599 |
| Average loop % | 43.11 |
| Average alpha helix % | 39.16 |
| Average beta sheet % | 17.74 |
| Average protein size | 338.93 |
| Mean Accuracy | 0.954 ± 0.036 |
| Mean Precision | 0.512 ± 0.135 |
| Mean Recall | 0.820 ± 0.141 |
| Mean Specificity | 0.959 ± 0.041 |
| Mean F1-score | 0.609 ± 0.095 |
| Mean MCC | 0.617 ± 0.086 |

For both datasets, available PDB structures were filtered by resolution (≤3.0 Å), and all NMR structures were retained to preserve conformational variability. For domains lacking experimental structures, AlphaFoldDB can be used as a complementary structural source.

### Contact Pattern Identification

Contact patterns were defined as spatial arrangements of up to five amino acid residues whose centers of mass are located within a maximum distance of 8.0 Å from each other. For each structure, all pairwise residue distances were computed, and clusters meeting this criterion were extracted as patterns. Each pattern was encoded using a simple notation: residues involved in contacts were represented by uppercase letters, and intervening residues by lowercase letters. Patterns were stored in a domain- or sequence-specific library for subsequent alignment.

#### Pattern Alignment and Contact Map Generation

Predicted contact maps were generated by aligning patterns from the library to query sequences using MATLAB’s(MATLAB R2023b) *localalign* function, with an increased gap-opening penalty of 10 to favor precise motif matching. When a pattern is successfully aligned to the query sequence, corresponding elements of the N × N contact matrix and their symmetric counterparts were incremented by 1. This process was repeated for all patterns in the domain or sequence-specific pattern library, cumulatively building the contact map through pattern alignment on the query sequence.

### Benchmarking and Validation

A two-tier benchmarking strategy was employed:

#### 1. Curated Domains (25 InterPro Domains)

For each of the 25 protein domains in our benchmark set (Table 1), we compared mean contact maps generated by our pattern matching approach with mean reference contact maps derived from experimentally determined structures.To address the substantial size difference between prediction datasets and experimental datasets, we employed a bootstrap resampling framework with 1,000 iterations. Multiple sequence alignments were generated using Clustal Omega^8^ to establish correspondence between predicted and experimental structures.

The statistical analysis implemented a dual-metric approach evaluating both full alignments and filtered regions, with Pearson correlation coefficients and bias metrics including mean absolute error and systematic deviation calculated across all aligned positions for full alignment analysis. In parallel, a filtering strategy addressed alignment quality differences by eliminating MSA columns with high gap content, calculating per-position coverage for both datasets and applying an intersection strategy to identify positions where both achieved at least 30% coverage for unbiased comparison. Statistical significance was established through bootstrap distributions enabling calculation of 95% confidence intervals and paired t-tests, with p-values computed for both comparison scenarios. The analysis extended to distance-dependent characterization examining sequence separations of 1, 5, 10, 15, 20, and 25 residues, and regional classification grouping contacts into short-range (|i-j| < 12), medium-range (12 ≤ |i-j| < 24), and long-range (|i-j| ≥ 24) categories. Performance evaluation encompassed comprehensive classification metrics including ROC curves with AUC values, precision-recall curves, and F1 scores at optimal thresholds, computed for both full and filtered analyses, providing statistically robust evidence for comparing datasets of vastly different sizes while accounting for varying data quality.

#### 2. Poorly Annotated Sequences (7,599 Unreviewed Proteins)

Predicted contact maps were validated against high-confidence AlphaFold models. Standard classification metrics were computed for each sequence, including accuracy, precision, recall, specificity, F1-score, and Matthews Correlation Coefficient (MCC). Population-level performance was summarized by mean and standard deviation across all sequences, and best/worst performers were identified to illustrate range of applicability.

This dual-validation framework allows robust assessment of method performance across datasets with vastly different structural coverage, ensuring applicability to both well-characterized protein families and poorly annotated sequences with limited structural information.

## Results

Application of the pattern-matching approach yielded contact maps that closely recapitulated the characteristic residue–residue networks of known protein folds. Across diverse protein families, the method reliably recovered conserved interaction patterns that could be directly compared with experimentally determined contact maps. This approach builds upon the successful framework demonstrated by Bradley, Kim, and Berger (2002)^9^ for identifying structural motifs that represent specific protein fold characteristics. As illustrated in Figure 1, this procedure reconstructs contact networks that align well with structural reference data.

**Figure 1:**
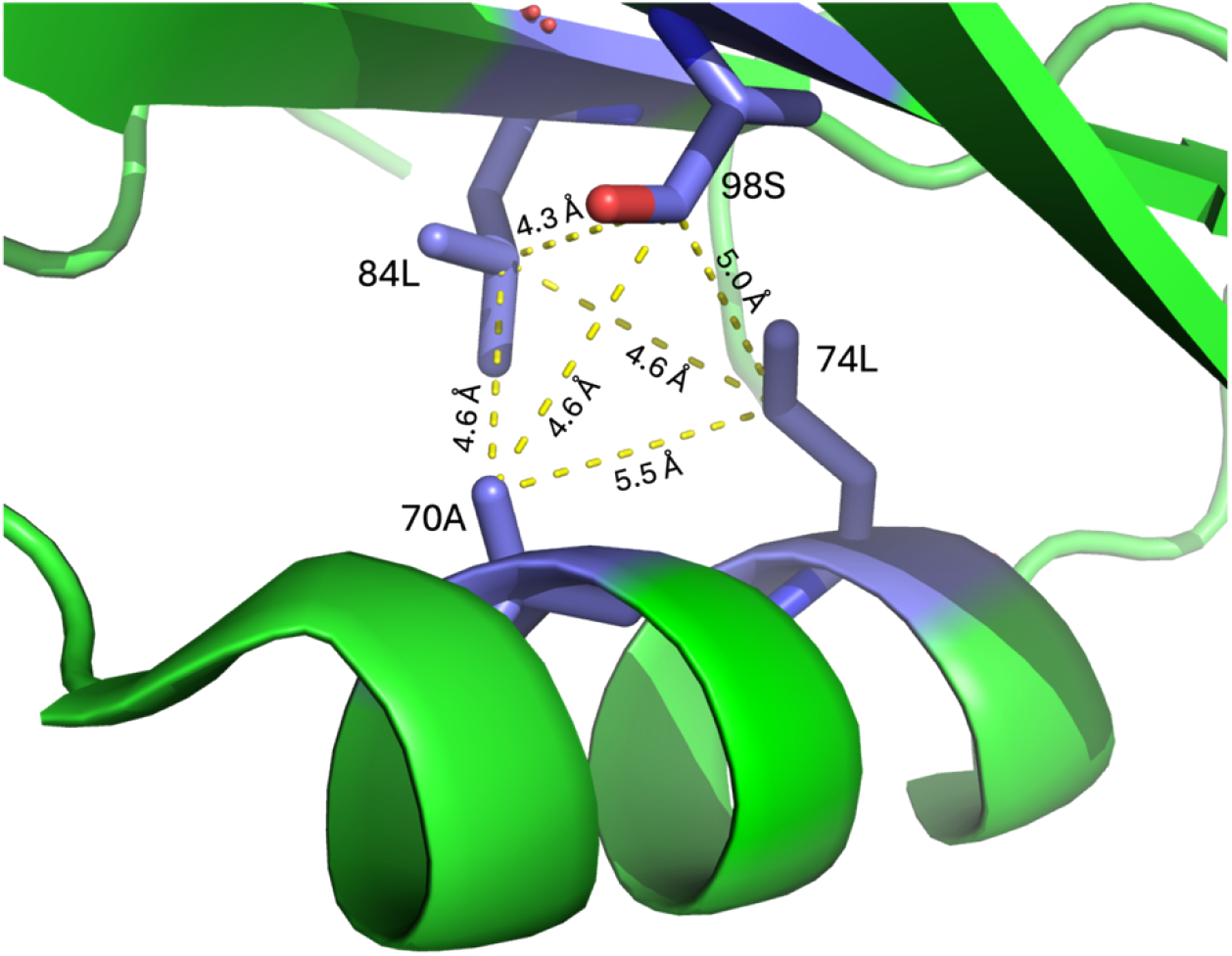
Example of a contact pattern in the Grb2 SH2 domain. Four residues (Ala70, Leu74, Leu84, and Ser98; shown as purple sticks) are located within 8.0 Å of each other, illustrating how spatially proximal residues are defined as a contact pattern (PDB ID: 6ICG)^10^). Contact patterns, defined as clusters of up to five residues within 8.0 Å, are aligned to query sequences. The example shows a sample pattern “**A**een**L**skqrhdgaf**L**ireresapgdfsl**S**” aligning against a query sequence from the human P56-LCK Tyrosine Kinase SH2 domain (PDB ID: 1LKK)^11^. Uppercase letters denote residues involved in contacts, lowercase letters represent intervening residues. The alignment begins at position 16 of the query sequence, with pattern residues A, L, L, and S corresponding to positions 16, 21, 31, and 45 in the query sequence. When a pattern aligns to a sequence, contacts are mapped to the corresponding positions in the contact matrix (C(16,21)=1, C(16,31)=1, C(16,45)=1, C(21,31)=1, C(21,45)=1, C(31,45)=1, along with symmetric counterparts). This process is repeated for all patterns in the library to generate the full predicted contact map.

To assess the performance of our approach, we applied it to two complementary datasets. The primary benchmark consisted of 25 well-characterized protein domains from InterPro^4^(Table 1). These domains are supported by well-curated sets of PDB structures, often capturing distinct conformational states, and thus provide a stringent test set with comprehensive structural annotation.

As a second benchmark, we assembled a dataset of sequences from UniProt’s^5^ TrEMBL^6^ collection. Unlike the curated InterPro domains, these proteins lack extensive experimental annotation. This dataset therefore evaluates the applicability of the method under conditions where structural information is sparse, reflecting a more realistic scenario for newly identified proteins.

### Performance on well-characterized protein domains

We first benchmarked our approach on 25 InterPro domains representing diverse architectures and supported by abundant structural data (Table 1). These domains span 20–250 residues and cover a wide range of secondary structure compositions, from loop-rich EGF-like domains (76.6% loops) to highly ordered spectrin repeats (9.0% loops). The large number of available experimental structures per domain (on average 473, up to 2,047 for the ubiquitin family) provided a robust basis for statistical evaluation.

Across all 25 families, the pattern-matching method showed robust performance, with correlation coefficients between predicted and reference contact maps ranging from 0.735 to 0.942. Classification accuracy was consistently high, with ROC AUC scores between 0.865 and 0.995, and F1 scores between 0.664 and 0.931. Thus, even in families with more modest correlations, discrimination between true and false contacts remained excellent (AUC >0.9 in all cases).

Family-specific differences were observed. Top-performing domains such as PF01023 (S-100), PF00505, PF00435 (Spectrin repeat), PF00001 (7TM receptor), and PF00249 consistently achieved correlations above 0.92 and F1 scores above 0.91, reflecting strong evolutionary constraints. By contrast, families such as PF00089 (Trypsin) showed lower correlations (0.735) but still maintained strong classification accuracy (AUC = 0.910).

Systematic biases in contact probability calibration varied by family, with small domains (25– 30 residues; e.g., PF00400, PF00096, PF00036) tending to under-predict contacts, while larger domains (83–250 residues; e.g., PF00089, PF00085, PF00017) showed slight over-prediction. These trends suggest that structural constraints and domain size jointly shape predictive behavior.

Contact range analysis revealed that short-range contacts (≤5 residues) were predicted most accurately (r = 0.772–0.929), while medium-range (5–15 residues) and long-range (>15 residues) contacts showed greater variability across families. Notably, the EF-hand domain (PF00036) achieved exceptional long-range prediction (r = 0.946), whereas the ligand-gated ion channel domain (PF00060) performed less well (r = 0.424).

Secondary structure composition strongly influenced predictive accuracy. Domains with higher loop content consistently exhibited reduced correlations: the loop-rich EGF-like domain (PF00008, 76.6% loops) reached r = 0.836, below the overall mean of 0.867. In contrast, the spectrin repeat (PF00435, 9.0% loops) achieved the highest correlation (r = 0.932). By contrast, domain size itself showed no significant correlation with accuracy, indicating that loop content, rather than length, is the primary determinant of performance.

Representative examples illustrate the range of outcomes. The S-100 domain (PF01023; r = 0.942, AUC = 0.972; Figure 2) exemplifies high accuracy, the RNA recognition motif (PF00076; r = 0.872, AUC = 0.988; Figure 3) shows intermediate performance, and trypsin (PF00089; r = 0.735, AUC = 0.911; Figure 4) represents lower correlations but still strong classification. Additional examples are provided in Figures S1–S50, with detailed metrics in Supplementary Text Files S1–S26.

**Figure 2:**
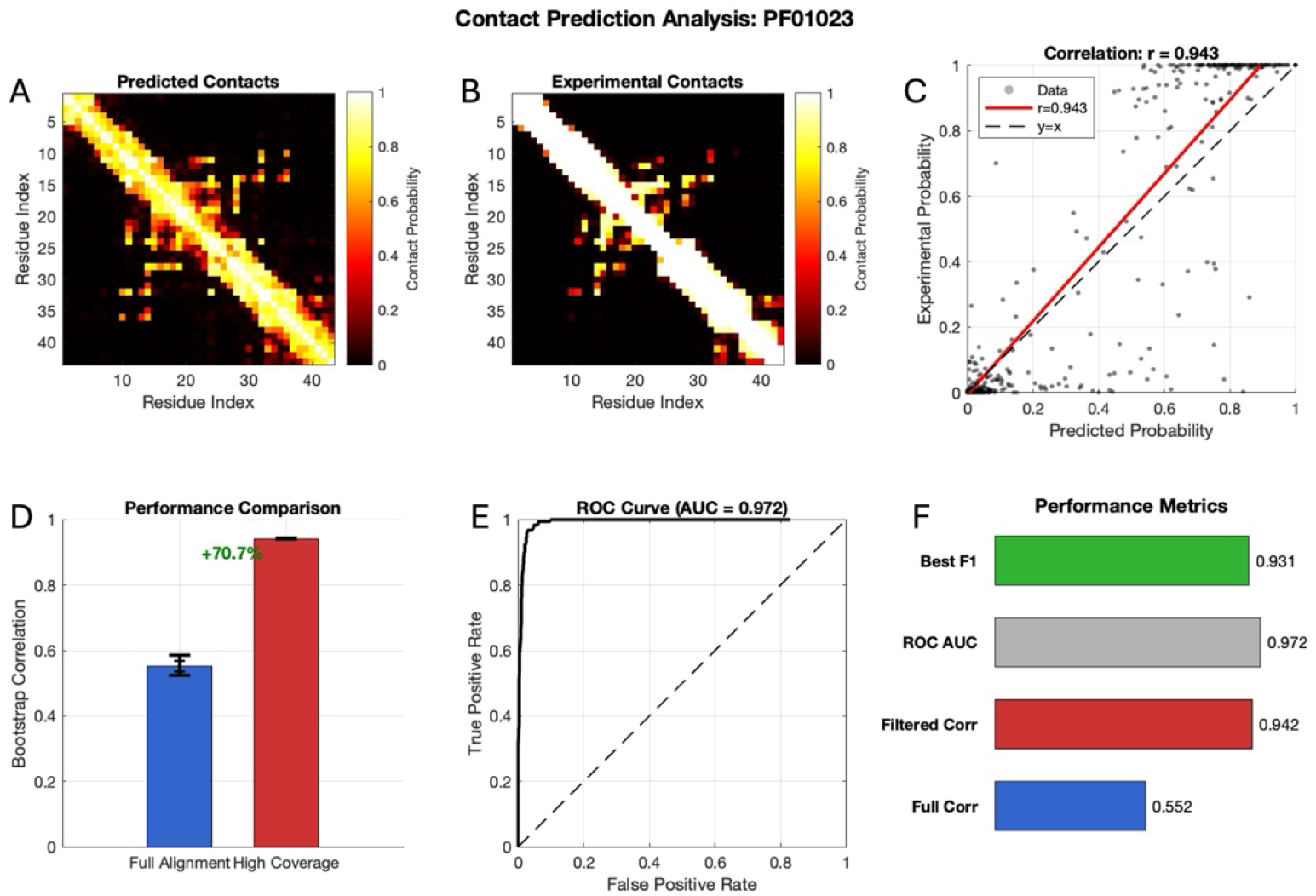
Example of a high-quality contact prediction for protein family PF01023 (S-100 domain). This case illustrates a very good prediction outcome. (A) Mean predicted contact map generated by our pattern-matching approach, showing contact probabilities between residue pairs (yellow: >0.8 probability; dark: low or no probability). (B) Experimental reference contact map derived from crystal structures for comparison. (C) Correlation analysis between predicted and experimental contact probabilities (r = 0.943). The red line shows the linear fit; the dashed black line indicates perfect correlation (y = x). (D) Comparison between full alignment analysis (blue, r = 0.552) and filtered high-coverage regions (red, r = 0.942), highlighting a 70.7% improvement with filtering. (E) ROC curve with area under the curve (AUC) of 0.972, indicating near-perfect discrimination between true and non-contacts. The dashed diagonal indicates random performance. (F) Summary of key metrics: best F1-score (0.931), ROC AUC (0.972), filtered correlation (0.943), and full alignment correlation (0.552).

**Figure 3:**
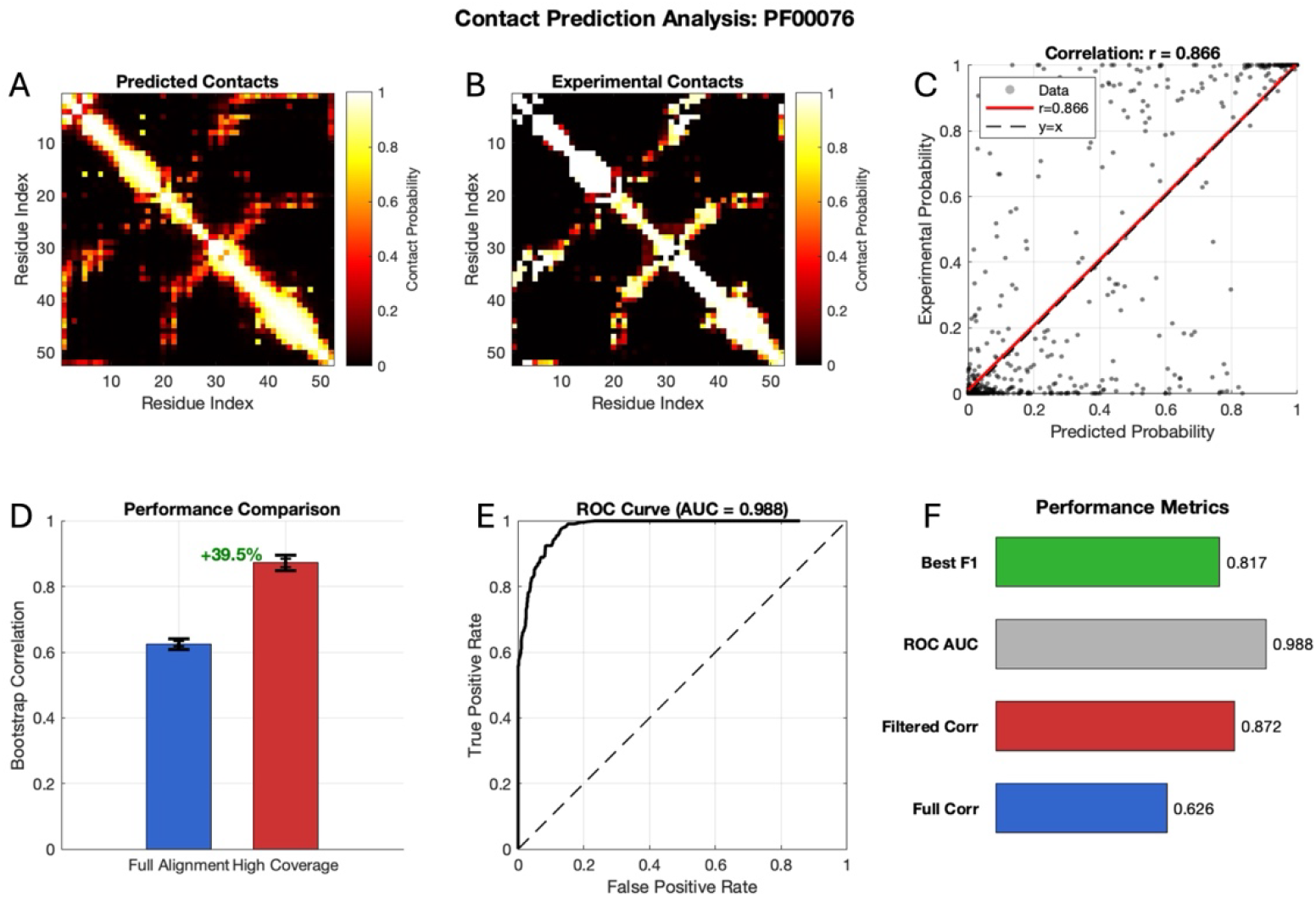
Example of a moderate contact prediction for protein family PF00076 (RNA recognition motif). This case illustrates a typical outcome for more challenging domains. (A) Predicted contact map from our pattern-matching approach, showing residue–residue contact probabilities (yellow/red: 0.4–0.8 probability; dark: low or no probability). (B) Experimental reference contact map derived from crystal structures, displaying a sparser contact distribution compared to well-conserved domains. (C) Correlation between predicted and experimental contact probabilities (r = 0.886). (D) Comparison of full alignment analysis (blue, r = 0.626) versus filtered high-coverage regions (red, r = 0.872), showing a 39.5% improvement with filtering. (E) ROC curve with area under the curve (AUC) of 0.988, indicating strong discriminative power even for this challenging family. (F) Summary of key metrics: best F1-score (0.817), ROC AUC (0.988), filtered correlation (0.872), and full alignment correlation (0.626).

**Figure 4:**
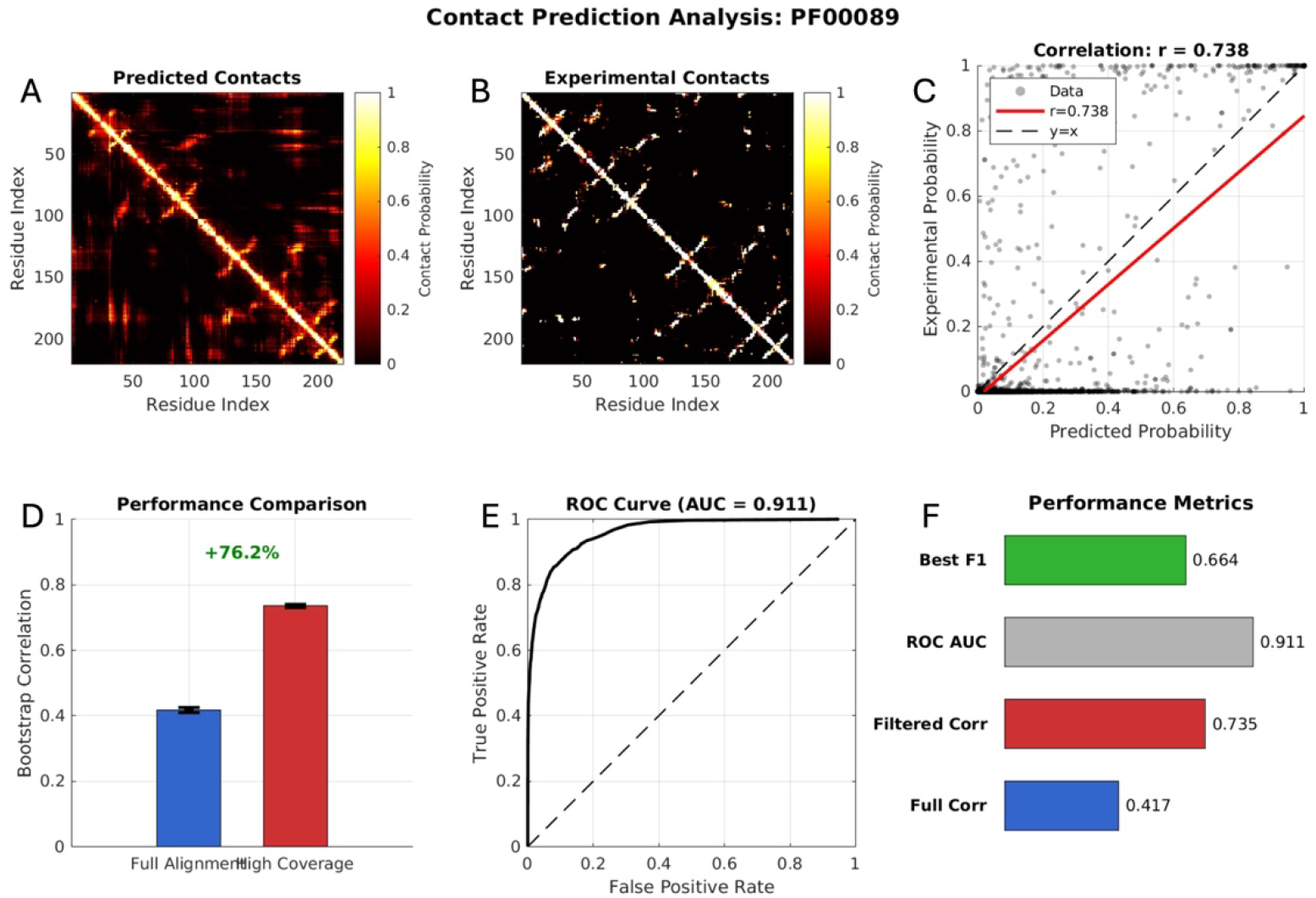
Example of a lower-quality contact prediction for protein family PF00089 (Trypsin). This case illustrates a poorer outcome, typical for highly diverse protein families. (A) Predicted contact map from our pattern-matching approach, showing residue–residue contact probabilities (yellow/red: 0.4–0.8; dark: low or none). (B) Experimental reference contact map derived from crystal structures. (C) Correlation between predicted and experimental contact probabilities (r = 0.738). The red line shows the linear fit; the dashed black line indicates perfect correlation (y = x). (D) Comparison of full alignment analysis (blue, r = 0.417) versus filtered high-coverage regions (red, r = 0.735), showing a 76.2% improvement with filtering. (E) ROC curve with area under the curve (AUC) of 0.911, indicating limited but still significant discriminative ability. The dashed diagonal indicates random performance. (F) Summary of key metrics: best F1-score (0.664), ROC AUC (0.911), filtered correlation (0.735), and full alignment correlation (0.417).

Overall, these results demonstrate that the pattern-matching approach achieves highly reliable contact prediction across structurally diverse protein families, with performance primarily influenced by secondary structure composition rather than domain length.

### Performance on poorly annotated sequences

To test the generalizability of our approach, we next evaluated 7,599 poorly annotated sequences, which lack curated experimental annotation and can only be represented by reference AlphaFold models. The method maintained robust predictive performance: over 90% of sequences achieved accuracies above 0.90 (Fig. 5A), and precision–recall analyses confirmed balanced classification with relatively few outliers (Fig. 5B–C). Additional measures, including specificity and Matthews Correlation Coefficient (MCC), consistently supported the reliability of contact predictions across diverse protein families (Fig. 5D–F).

**Figure 5:**
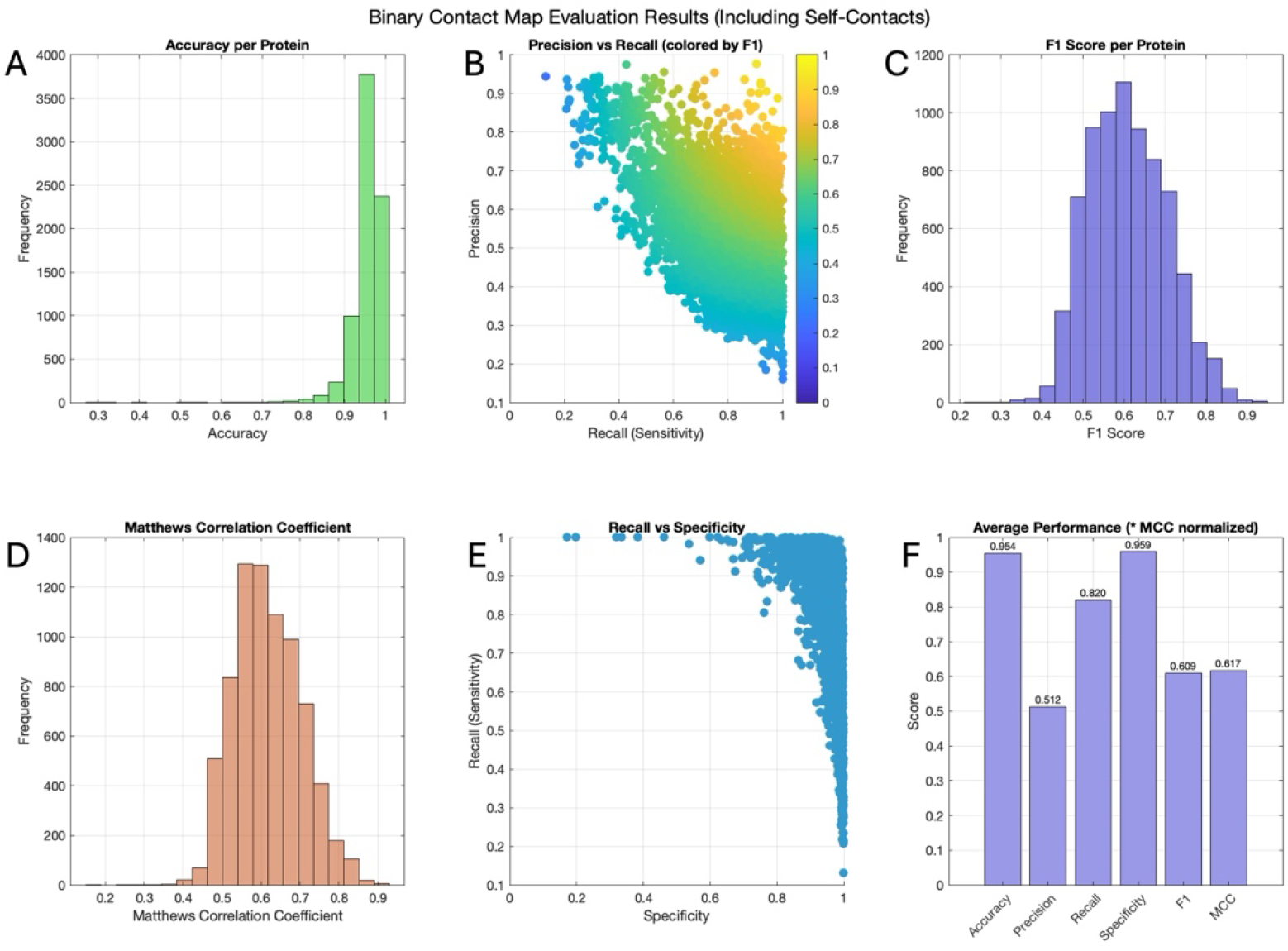
Comprehensive binary classification performance analysis for 7,599 poorly annotated sequences using homologous structural templates. (A) Accuracy distribution across all analyzed sequences, showing consistently high performance with most sequences achieving >95% accuracy. The narrow distribution (mean: 0.954 ± 0.036) demonstrates robust classification performance across diverse protein families and sizes (20-2,017 residues). (B) Precision-recall relationship colored by F1-score, revealing the method’s characteristic high sensitivity (recall) with moderate precision. The color gradient from blue to yellow indicates F1-scores ranging from 0.10 to 0.90, with the majority of sequences achieving F1-scores between 0.5-0.8. The broad recall range (0.2-1.0) reflects varying contact density across different protein architectures. (C) F1-score distribution showing a right-skewed distribution with mean performance of 0.609 ± 0.095, indicating that most sequences achieve balanced precision-recall performance with relatively few poorly performing outliers. (D) Matthews Correlation Coefficient (MCC) distribution demonstrating substantial agreement between predicted and AlphaFold reference contact maps, with mean MCC of 0.617 ± 0.086. The distribution peak around 0.6-0.7 indicates reliable binary classification performance across the dataset. (E) Recall versus specificity analysis showing the method’s high true positive rate (mean recall: 0.820) combined with excellent specificity (mean: 0.959), indicating effective identification of true contacts with minimal false positive rates. The concentrated distribution in the upper-right quadrant demonstrates consistent performance across diverse protein families. (F) Summary of average performance metrics across all sequences, with MCC values normalized for comparison. The high specificity (0.959) and accuracy (0.954) demonstrate the method’s reliability for practical contact prediction applications, while the moderate precision (0.512) reflects the challenging nature of contact prediction for diverse protein families using homologous templates. The balanced F1-score (0.609) and substantial MCC (0.617) indicate meaningful structural information extraction across the entire dataset of poorly annotated sequences.

Prediction accuracy was strongly affected by secondary structure content. Proteins dominated by well-ordered helices or sheets showed near-perfect predictions—for example, A2AIM4, with only 1% loop residues, achieved accuracy = 0.995 and MCC = 0.919. By contrast, highly disordered proteins performed more poorly: Q7M4S4, composed entirely of loops, reached only 0.550 accuracy and MCC = 0.379. Representative examples of high-, medium-, and low-accuracy cases are shown in Figures 6–8. Overall, these results highlight that loop-rich or intrinsically disordered regions are the main challenge for contact prediction, whereas structured domains yield excellent agreement with reference models.

**Figure 6:**
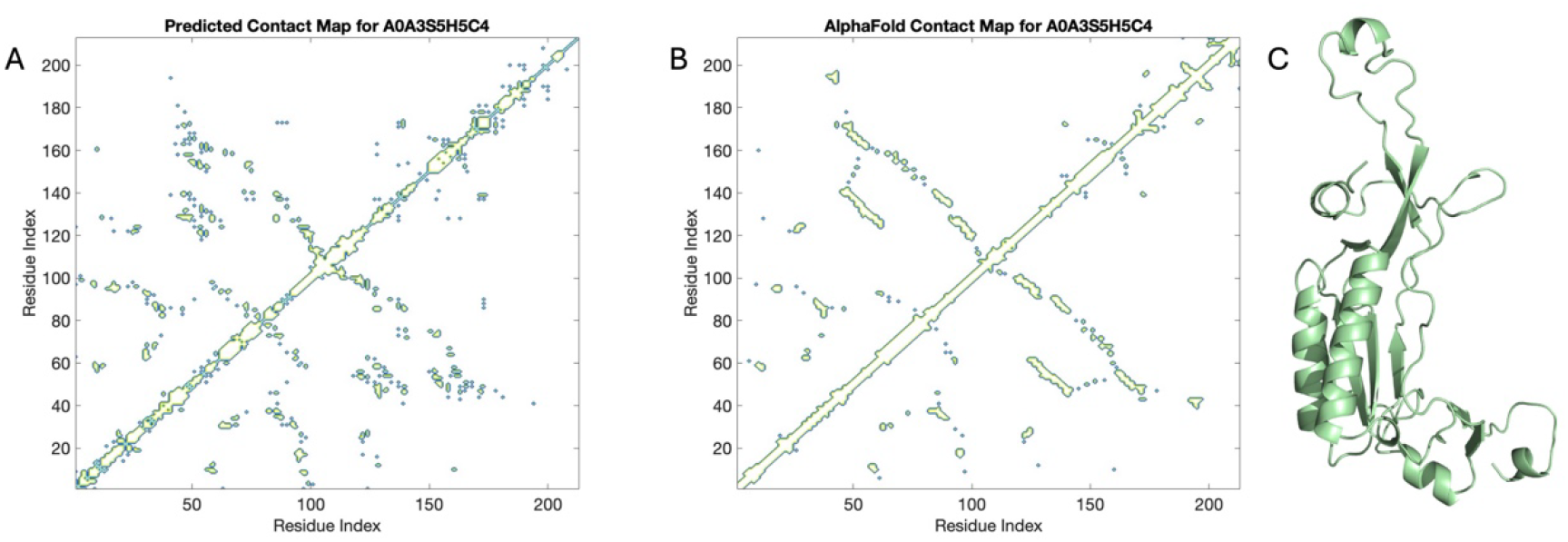
Comparison of predicted and AlphaFold contact maps for protein A0A3S5H5C4 (60S ribosomal protein L10 from Leishmania donovani). (A) Predicted contact map. (B) Reference contact map derived from the AlphaFold structure. The prediction shows high accuracy (96.8%) and specificity (98.8%), with moderate precision (69.8%) and recall (57.2%), yielding an F1-score of 0.629 and MCC of 0.615. Performance corresponds to 1,239 true positives, 536 false positives, 42,666 true negatives, and 928 false negatives out of 45,369 residue pairs. (C) AlphaFold structure shown in cartoon representation; the loop content of this protein is 50.7%.

**Figure 7:**
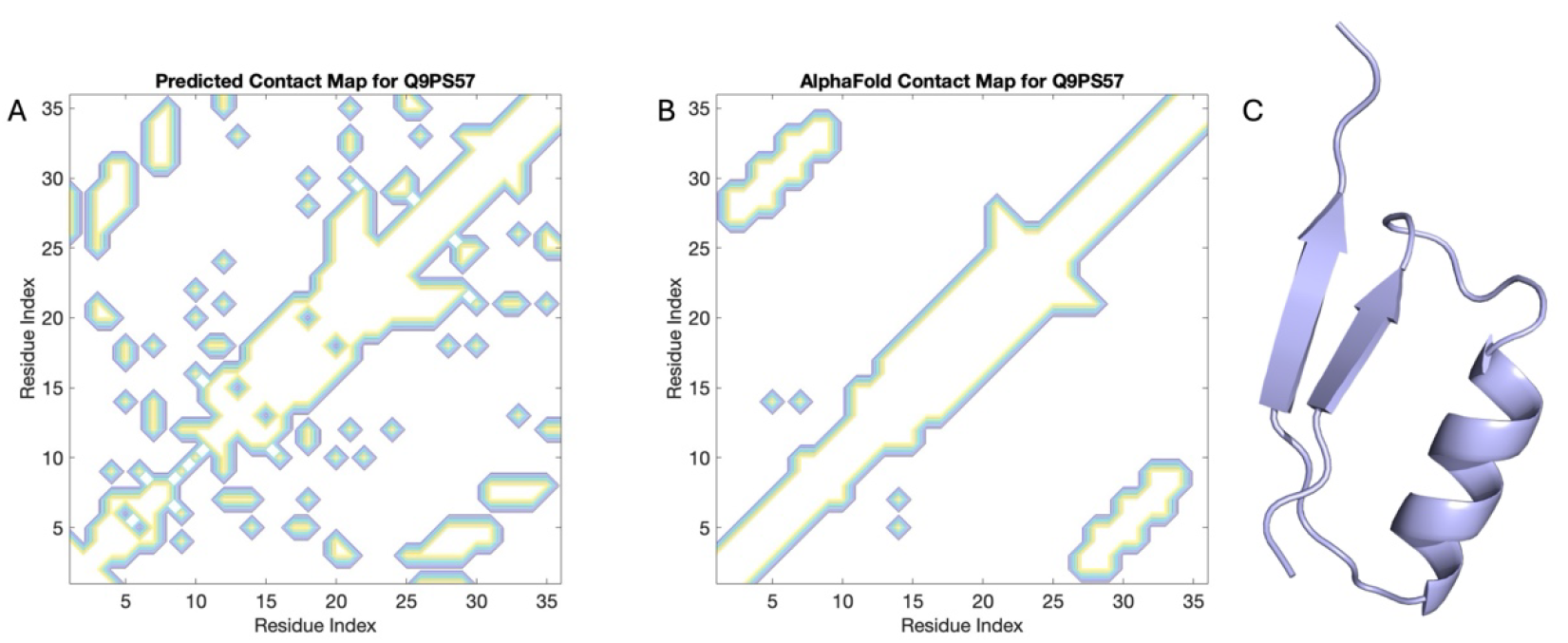
Comparison of predicted and AlphaFold contact maps for protein Q9PS57 (Glutathione transferase isoenzyme III from Bufo bufo). (A) Predicted contact map. (B) Reference contact map derived from the AlphaFold structure. The prediction shows an accuracy of 87.5% and specificity of 89.2%, with moderate precision (68.8%) and recall (81.5%), yielding an F1-score of 0.746 and MCC of 0.668. Performance corresponds to 238 true positives, 108 false positives, 896 true negatives, and 54 false negatives out of 1,296 residue pairs. (C) AlphaFold structure shown in cartoon representation; the loop content of this protein is 44.4%.

**Figure 8:**
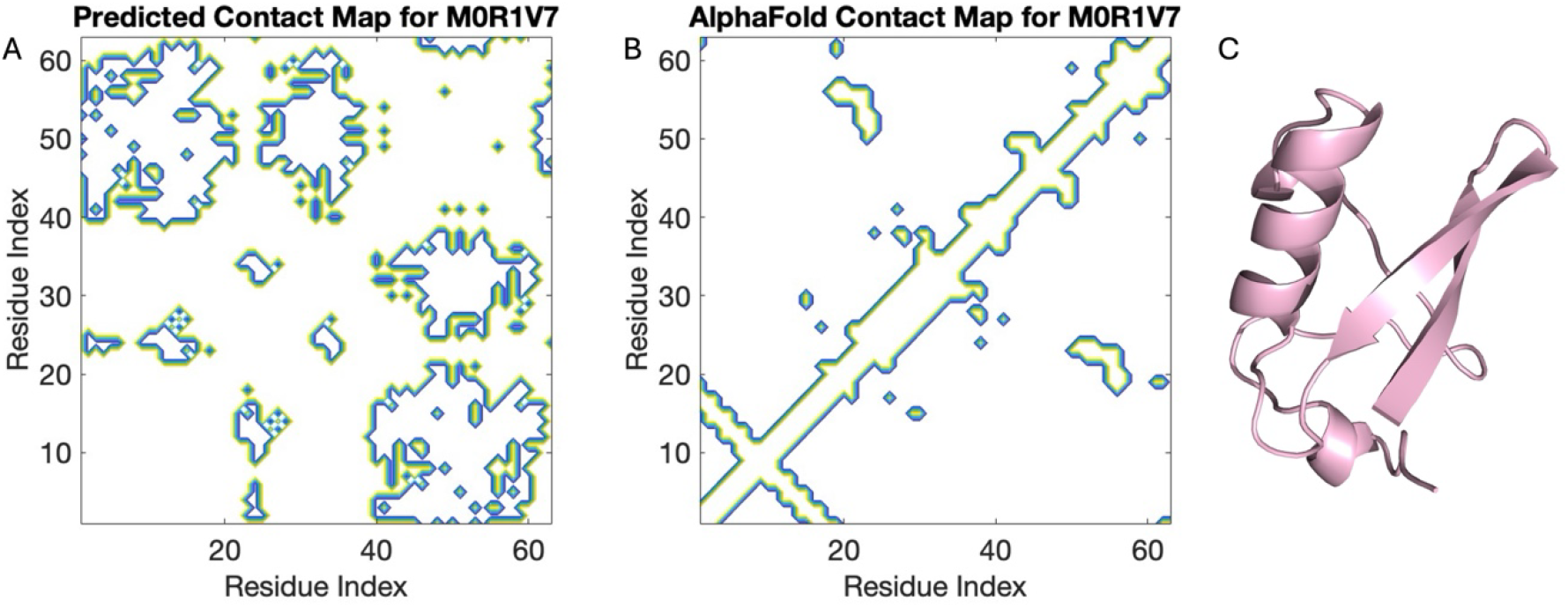
Comparison of predicted and AlphaFold contact maps for protein M0R1V7 (Ubiquitin A-52 residue ribosomal protein fusion product 1 from Homo sapiens). (A) Predicted contact map. (B) Reference contact map derived from the AlphaFold structure. The prediction shows an accuracy of 41.7% and specificity of 32.0%, with low precision (19.6%) and recall (1.0%), yielding an F1-score of 0.328 and MCC of 0.251. Performance corresponds to 565 true positives, 2,314 false positives, 1,090 true negatives, and 0 false negatives out of 3,969 residue pairs. (C) AlphaFold structure shown in cartoon representation; the loop content of this protein is 46.0%.

We also examined whether the number of structural homologs in the PDB influences prediction accuracy. Surprisingly, performance showed a negative correlation on the number of PDB hit counts with MCC and F1 scores, the correlation values are -0.164 and -0.178 respectively; Fig. 9B– C). Grouping sequences by homolog abundance revealed slightly declining MCC and F1 scores as hit counts increased (Fig. 9D). Thus, adding more distant homologs does not improve performance and may in fact dilute the signal. Tables 3 and 4 list the best and worst performers, further illustrating that accuracy is not driven by homolog availability.

**Table 3.** Top 10 sequences ranked by MCC and F1-score

| Top 10 performers | MCC | F1-score | Accuracy | PDB Hit count |
| --- | --- | --- | --- | --- |
| A0A151DYG7 | 0.927 | 0.939 | 0.978 | 83 |
| A7XZE4 | 0.923 | 0.925 | 0.995 | 62 |
| A0A4X1VW25 | 0.92 | 0.923 | 0.995 | 62 |
| A2AIM4 | 0.919 | 0.922 | 0.995 | 62 |
| A0A1Y7VNT9 | 0.911 | 0.933 | 0.964 | 288 |
| D0VX28 | 0.895 | 0.907 | 0.978 | 68 |
| Q6CM20 | 0.892 | 0.897 | 0.99 | 249 |
| A0A140LIU3 | 0.886 | 0.906 | 0.963 | 68 |
| Q6CPN9 | 0.885 | 0.893 | 0.985 | 241 |
| A0A804RMZ3 | 0.878 | 0.881 | 0.992 | 89 |

**Table 4.** Bottom 10 sequences ranked by MCC and F1-score

| Bottom 10 Performers | MCC | F1-score | Accuracy | PDB Hit count |
| --- | --- | --- | --- | --- |
| Q9PST8 | 0.166 | 0.277 | 0.285 | 890 |
| F5GZ39 | 0.186 | 0.297 | 0.314 | 872 |
| M0R1V7 | 0.251 | 0.328 | 0.417 | 881 |
| A0A0P0W7F3 | 0.299 | 0.31 | 0.606 | 807 |
| C4M760 | 0.309 | 0.343 | 0.528 | 898 |
| Q9UMG3 | 0.346 | 0.528 | 0.515 | 177 |
| A0A498MMR3 | 0.351 | 0.232 | 0.991 | 201 |
| Q0JBH4 | 0.365 | 0.364 | 0.695 | 856 |
| D3YVJ8 | 0.372 | 0.36 | 0.716 | 894 |
| Q7M4S4 | 0.379 | 0.545 | 0.55 | 612 |

**Figure 9:**
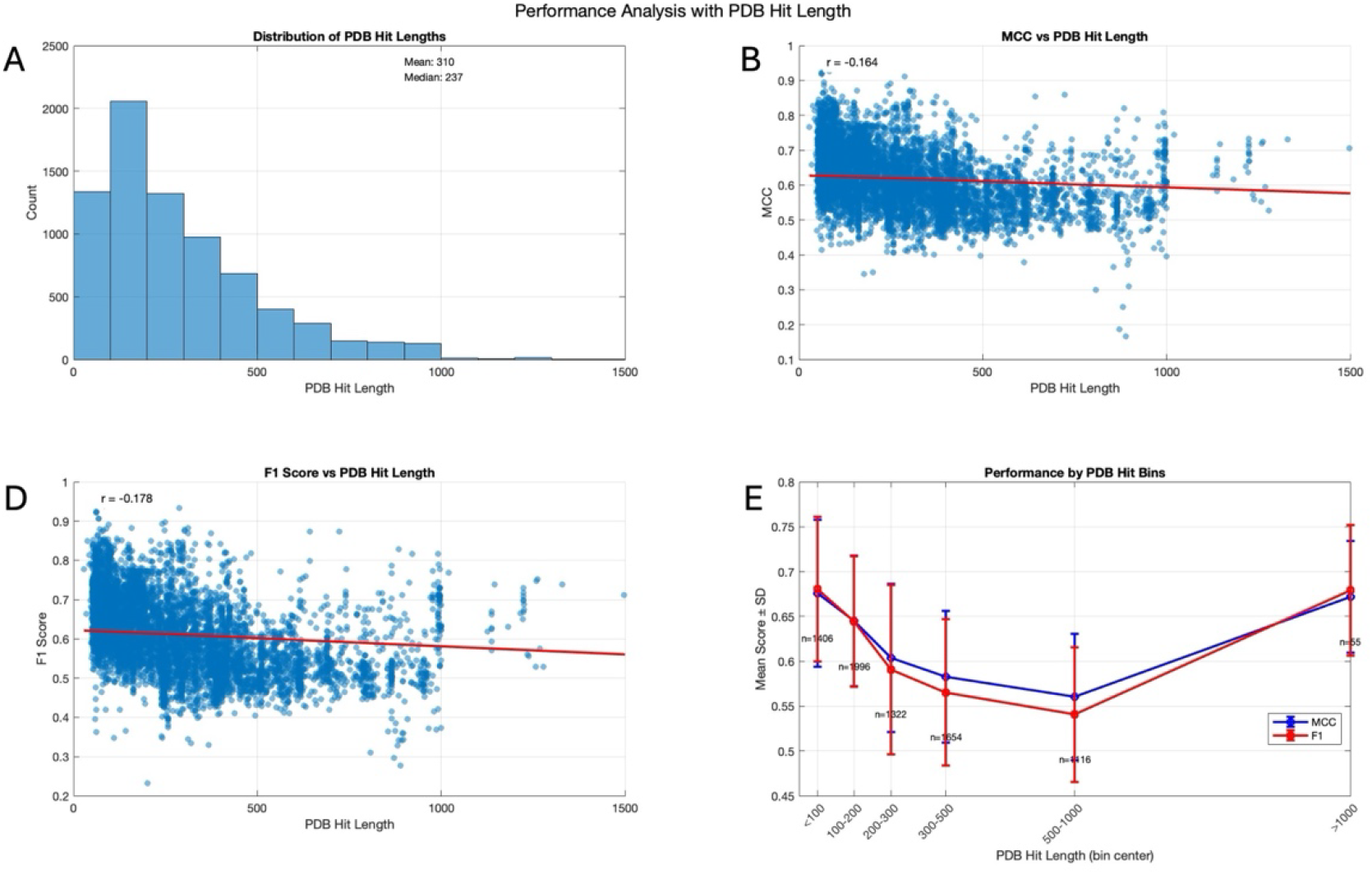
Performance Analysis of Protein Function Prediction Model Relative to PDB Hit Lengths. (A) Distribution of PDB hit counts. (B) Scatter plot revealing weak negative correlation (r = -0.164) between PDB hit count and MCC. (C) Similar negative correlation (r = -0.178) between PDB hit count and F1-score. (D) Mean MCC values and F1-scores are grouped by PDB hit counts.

These findings demonstrate that the pattern-matching approach remains effective for poorly annotated proteins, even when only predicted structures are available. Performance is primarily determined by secondary structure composition rather than dataset curation or homolog abundance. The comparable results between curated domains and unreviewed sequences suggest that the essential structural motifs captured by the 8.0 Å contact patterns are conserved across wide evolutionary distances, supporting the biological relevance of this representation for newly discovered proteins, including archaeal candidates.

## Discussion

Understanding the protein structure is essential for explaining the biological function, yet the gap between sequenced proteins and experimentally determined structures continues to widen. In this study, we developed a pattern-matching approach that uses existing structural data to rapidly generate contact maps from sequence information alone. By validating the method on both well-characterized protein domains and poorly annotated sequences, we demonstrate its versatility, robustness, and practical utility across a wide range of proteins.

Our approach accurately predicts contacts across diverse protein families, capturing both short- and long-range interactions. Prediction performance is influenced by protein architecture, secondary structure composition, and evolutionary constraints, with well-structured domains yielding the most reliable results and loop-rich or intrinsically disordered proteins remain more challenging, highlighting the method’s dependence on conserved structural motifs for optimal accuracy.

The method also performs robustly for poorly annotated sequences, using distant homologs to generate meaningful contact predictions. This demonstrates its applicability to metagenomic datasets and proteins from organisms with limited structural coverage. Examples of such uncharacterized proteins include archaeal sequences, where our predictions show good agreement with AlphaFold reference structures (supplementary contact maps provided). Interestingly, increasing the number of structural homologs does not necessarily improve prediction quality, suggesting that a modest set of representative templates are sufficient to capture essential structural patterns.

A key advantage of our approach is its computational efficiency and interpretability, requiring minimal resources while revealing which structural motifs drive contact predictions. By integrating patterns from multiple structural states, the method inherently captures conformational flexibility, providing insights into dynamic protein behavior and allosteric mechanisms that would otherwise require extensive molecular dynamics simulations. This capability enables practical applications in functional annotation and drug discovery, allowing researchers to identify potential binding sites, infer allosteric pathways with additional methods, and guide structural characterization for novel proteins, even when experimental structures are unavailable.

Despite these advantages, the approach has limitations. Prediction accuracy is reduced for proteins dominated by loops or disorder, and the current 8.0 Å contact cutoff may not capture all functionally relevant long-range interactions in very large proteins. Future improvements could involve adaptive cutoffs or expanded pattern libraries to enhance coverage and accuracy.

Our study demonstrates that pattern-based contact map prediction provides a practical alternative to full structure prediction, bridging the gap between sequence and structure in a computationally efficient manner. As structural databases continue to expand, this approach can be readily adapted for exploratory studies in emerging protein families.

## Abbreviations

GPU: graphics processing unit
MSA: multiple sequence alignment
ROC: Receiver Operating Characteristic
AUC: Area Under the curve
MCC: Matthews Correlation Coefficient
pLDDT: predicted Local Distance Difference Test

## Availability of the code

The data that support the findings of this study are openly available in the Zenodo repository with the link https://zenodo.org/records/17043595.

Acknowledgments

This work was supported by the Swiss National Science Foundation (Grant 310030_219549 to D.F.). We thank the Division de Calcul et Soutien à la Recherche of the UNIL for access to the university’s computer infrastructure. We thank all members of the Fasshauer Laboratory for helpful discussions.

## Author contributions

A.H. designed the study, performed the experiments, and analyzed the data; A.H. and D.F. wrote the paper.

